# Downregulation of NCL attenuates tumor formation and growth in cervical cancer by targeting the PI3K/AKT pathway

**DOI:** 10.1101/2021.04.01.438118

**Authors:** Jun Ying, Ruowang Pan, Zhouhao Tang, Jiayin Zhu, Ping Ren, Yang Lou, Enyong Zhang, Dadao Huang, Penghong Hu, Dong Li, Qiyu Bao, Peizhen Li

## Abstract

Nucleolin (NCL, C23) is a multifunctional phosphoprotein that plays a vital role in modulating the survival, proliferation and apoptosis of cancer cells. However, the effects of NCL on cervical cancer and the underlying mechanisms behind this are poorly understood. In the study presented here, Hela cells were transfected with shRNAs targeting the endogenous NCL gene (sh-NCL-Hela). NCL knockdown inhibited cell proliferation and promoted apoptosis both *in vivo* and *in vitro*. Mechanistic studies revealed that NCL knockdown inhibited the PI3K/AKT pathway by upregulating FGF, ITGA, TNXB, VEGF, Caspase 3, and Bax, as well as by downregulating AKT, GNB4, CDK6, IL6R, LAMA, PDGFD, PPP2RSA and BCL-2. In addition, the expression levels of apoptosis-related genes after using a PI3K inhibitor LY294002 were consistent with shRNA studies, while treatment with a 740Y-P agonist showed the opposite effect. Altogether, this study uncovered that downregulation of NCL may be a novel treatment strategy for cervical cancer.

## Introduction

Cervical cancer is a common malignant tumor of the female reproductive system with an increasing incidence (1). There are about 500,000 new cervical cancer cases worldwide each year, of with more than 130,000 cases being diagnosed in China (2). According to the World Health Organization (WHO), about one million patients worldwide will die of cervical cancer each year by the year 2050 based on the projected rate. In China, 20,000 to 30,000 people die from cervical cancer each year, even at a younger age. In addition, the 5-year survival rate for cervical cancer is only 45.4% (3). Despite recent advances in cancer diagnosis and therapy, the prognosis and quality of life of patients are still not optimistic. Therefore, discovering new molecular targets and markers, increasing the survival rate, improving the prognosis and curative effects and achieving precise treatment need to be the priority of research in the future.

In our previous work, the expression levels of nucleolin (NCL) in cervical cancer cells were shown to be downregulated after treatment with phycocyanin (PC) as shown by two-dimensional gel electrophoresis (2-DE) and mass spectrometry (MALDI-TOF/MS), suggesting that NCL plays an important role in the development, cell proliferation and apoptosis pathways. Studies have showed that NCL exhibits multiple biological functions, including regulation of cell proliferation and growth, embryogenesis, cytokinesis, chromatin replication and the occurrence of nucleolus and other processes (4, 5), It plays a regulatory role by directly binding to related proteins in during DNA replication, recombination, and repair. Interestingly, NCL has recently been identified as an apoptosis-related proteins. Otake *et al*. found that Bcl-2 protein levels increased 11-fold when NCL increased 26 fold in B-lymphocytic leukemia (6). The expression levels of 110 kDa NCL in the nucleus reduced but an 80 kDa NCL degradation fragment appeared in human salivary gland cells (HSG), human oral squamous cell carcinoma cells (SCC-25) and human osteoblasts (Saos-2 and MG63) when induced by okadaic acid (7, 8). In addition, Sanjeev *et al*. conducted miRNA transcriptomics analysis using HeLa cells depleted in NCL and found that the down-regulation of NCL was associated with a decrease in cholesterol levels and an increase in fatty acid content, which was mainly due to the decreased and mis-localized expression of the transcription factor SREBP1 and the down-regulation of enzymes involved in β-oxidation and degradation of fatty acids (9). The expression levels of NCL are relatively high in tumors and other rapidly dividing cells, but very low in non-dividing cells. Using this feature as an effective indicator, NCL may be used to measure the degree of tumor cell proliferation and to judge sensitivity to certain drugs.

Even though there is accumulating evidence proving the pro-tumor activity of NCL, data are still limited and the potential molecular mechanisms remain unknown. A treatment strategy using NCL as an entry point will bring hope to personalized treatment of diseases, especially tumors. In this study, we systematically investigated the role and underlying mechanisms of NCL knockdown in relation to cell growth both *in vitro* and *in vivo* to provide more detailed theoretical data and an experimental basis for the functional mechanisms of NCL and the diagnosis and treatment of cervical cancer.

## Results

### Knockdown of NCL using shRNAs

The efficiency of NCL knockdown in Hela cells was verified by analyzing mRNA and protein expression levels. Compared with sh-NT-Hela cells, NCL mRNA and protein expression levels in sh-NCL-Hela were reduced compared with sh-NT-Hela cells (RT-PCR: t=18.048, ^*^*P* < 0.05; WB: t=14.008, ^**^*P* < 0.01), while there were no obvious differences observed between the sh-NT-Hela and NC-Hela cell lines (Figure 1A-B). The expression levels of β-actin mRNA and protein were similar in all groups. This suggested that NCL was significantly downregulated in sh-NCL-Hela cells.

**Figure 1:**
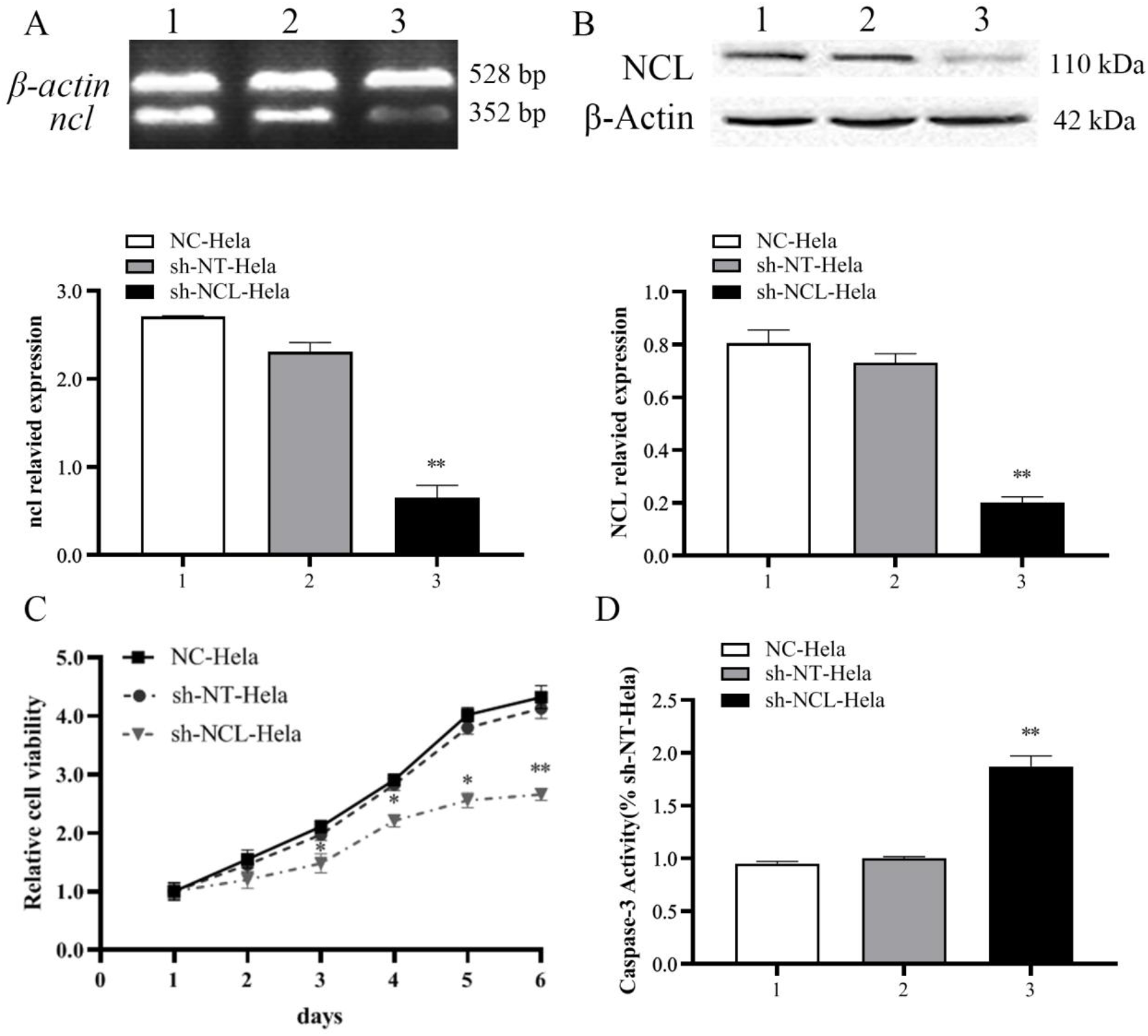
RNA interference inhibited NCL expression levels in HeLa cells. **A.** NCL mRNA expression detected by RT-PCR. **B.** NCL protein expression detected by Western blotting. **C.** Cell proliferation detected by MTT analysis at 1-, 2-, 3-, 4-, 5- and 6 days. **D.** Caspase-3 activity. Experiments were repeated three times. ^*^*P* < 0.05, ^**^*P* < 0.01, vs. sh-NT-Hela group cells, *n*=3.

### NCL knockdown inhibits cell proliferation in vitro

Compared with sh-NT-Hela cells, sh-NCL-Hela proliferation was notably inhibited in a time-dependent manner, and the highest inhibitory rate was 40.0 ± 1.7% on day 6. However, there were no obvious differences observed in the sh-NT-Hela and NC -Hela lines (Figure 1C). Colony formation assays revealed fewer and smaller colonies in sh-NCL-Hela cells than in sh-NT-Hela and NC -Hela cells (^**^*P* < 0.01) (Figure 2A). These results indicated that knockdown of NCL expression significantly inhibited the growth of Hela cells *in vitro*.

**Figure 2:**
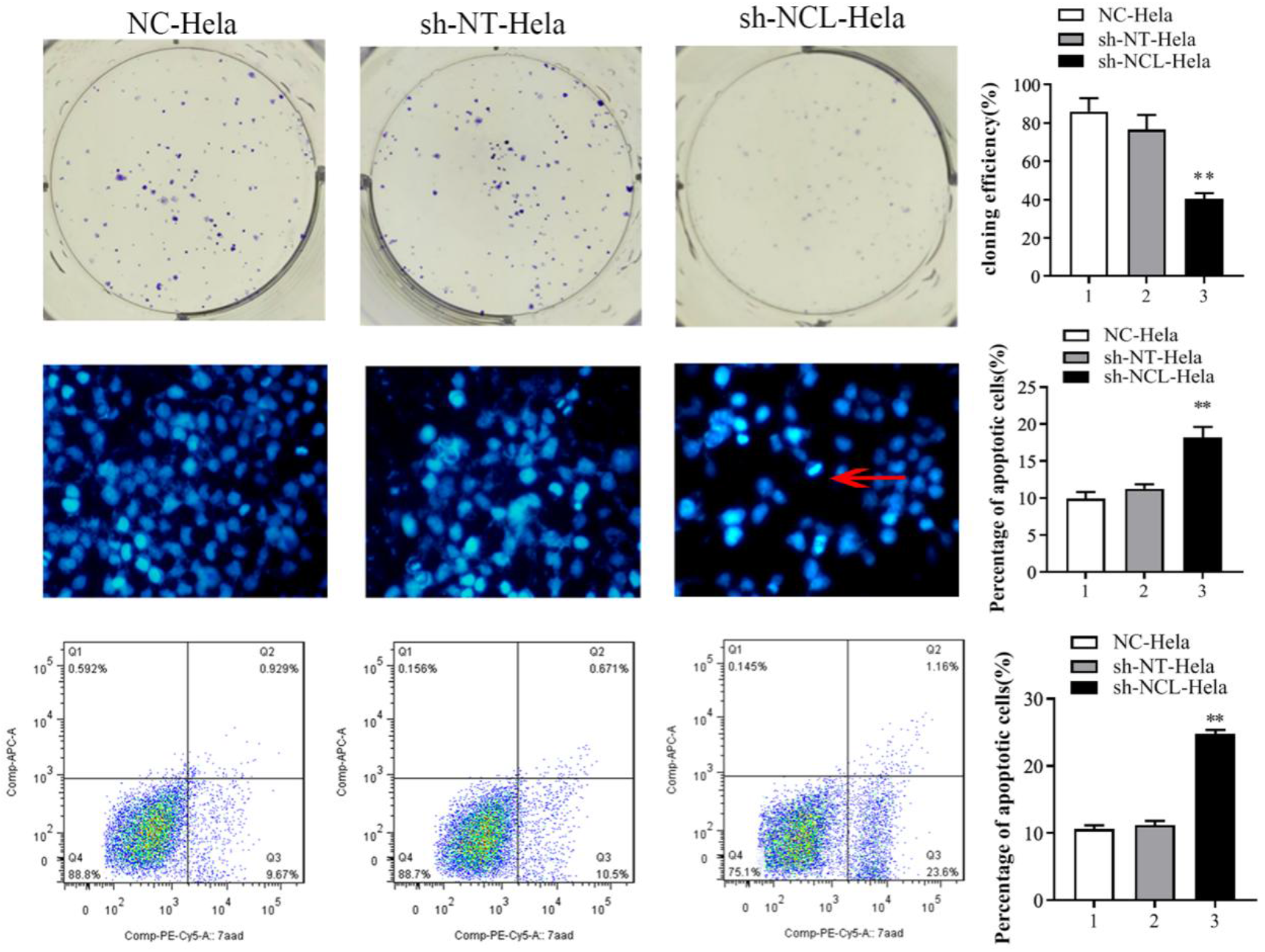
Detection of proliferation and apoptosis of Hela cells after NCL interference. **A.** Colony formation assay. Cells were inoculated into plates at a density of 500 cells per well and grown at 37°C for 14 days. Cell colonies were stained with 0.1% crystal violet (Left). Cell colonies were quantified, ^**^*P* < 0.01 (Right). **B.** NCL knockdown induced apoptosis morphological features as shown by Hoechst. Hypercoagulable nuclei and semilunar apoptotic bodies represent typical apoptosis morphology (see the red arrow; fluorescence staining × 400). **C.** Cell apoptosis was detected using flow cytometry. Representative flow cytometry analysis of Annexin V-APC/7AAD staining (Left). Apoptosis rate was quantified (Right), ^**^*P* < 0.01, vs. sh-NT-Hela group cells, *n*=3.

### NCL knockdown promotes apoptosis in Hela cells

Hoechst 33342 staining was performed to identify how NCL knockdown regulated cell death. Results showed that there was a large number of hypercoagulable nuclei and semilunar apoptotic bodies in sh-NCL-Hela cells (Figure 2B) and that apoptosis was more obvious at 48 h versus 24 h. Then we confirmed these finding using flow cytometry following Annexin V-APC/7AAD staining. Consequently, a significant increase in apoptotic cells was observed in sh-NCL-Hela cells (24.76 ± 0.59%), while low levels were observed in sh-NT-Hela cells (11.17 ± 0.63%) and NC-Hela cells (10.60±0.53%). There were significant differences noted between sh-NCL-Hela cells and sh-NT-Hela cells (^**^*P* < 0.01), no differences observed between sh-NT-Hela cells and NC -Hela cells (*P*>0.05) (Figure 2C). Similarly, knockdown of NCL also increased Caspase-3 enzyme activity (Figure 1D).

### NCL knockdown significantly inhibited tumor formation of Hela in vivo

To examine the effects of NCL knockdown on *in vivo* tumor growth, Hela cells were inoculated in nude mice and tumor volumes were measured up to 28 days. As shown in Figure 3, mice inoculated with sh-NCL-Hela cells showed a significantly lower tumor growth curve (^*^*P* < 0.05). At the terminal period, the mean tumor volume in the sh-NCL-Hela group was 0.38 times smaller than the sh-NT-Hela group (626.9 ± 263.8cm^3^ vs.1614 ± 272.6 cm^3^, ^*^*P* < 0.05), with an average weight being 0.513 ±0.18 g for sh-NCL-Hela group and 1.454 ± 0.38 g for sh-NT-Hela group (^*^*P*<0.05). Using H&E staining, the degree of cell differentiation was higher in sh-NCL-Hela group compared to the control group and the nucleus to cytoplasm ratio was also significantly reduced. Furthermore, TUNEL staining showed that the number of apoptosis-positive cells in the sh-NCL-Hela group (99.7 ± 12.6%) was significantly increased compared to the sh-NT-Hela control group (0.9 ± 0.73%). Altogether, NCL knockdown in Hela cells markedly suppressed *in vivo* tumor growth.

**Figure 3:**
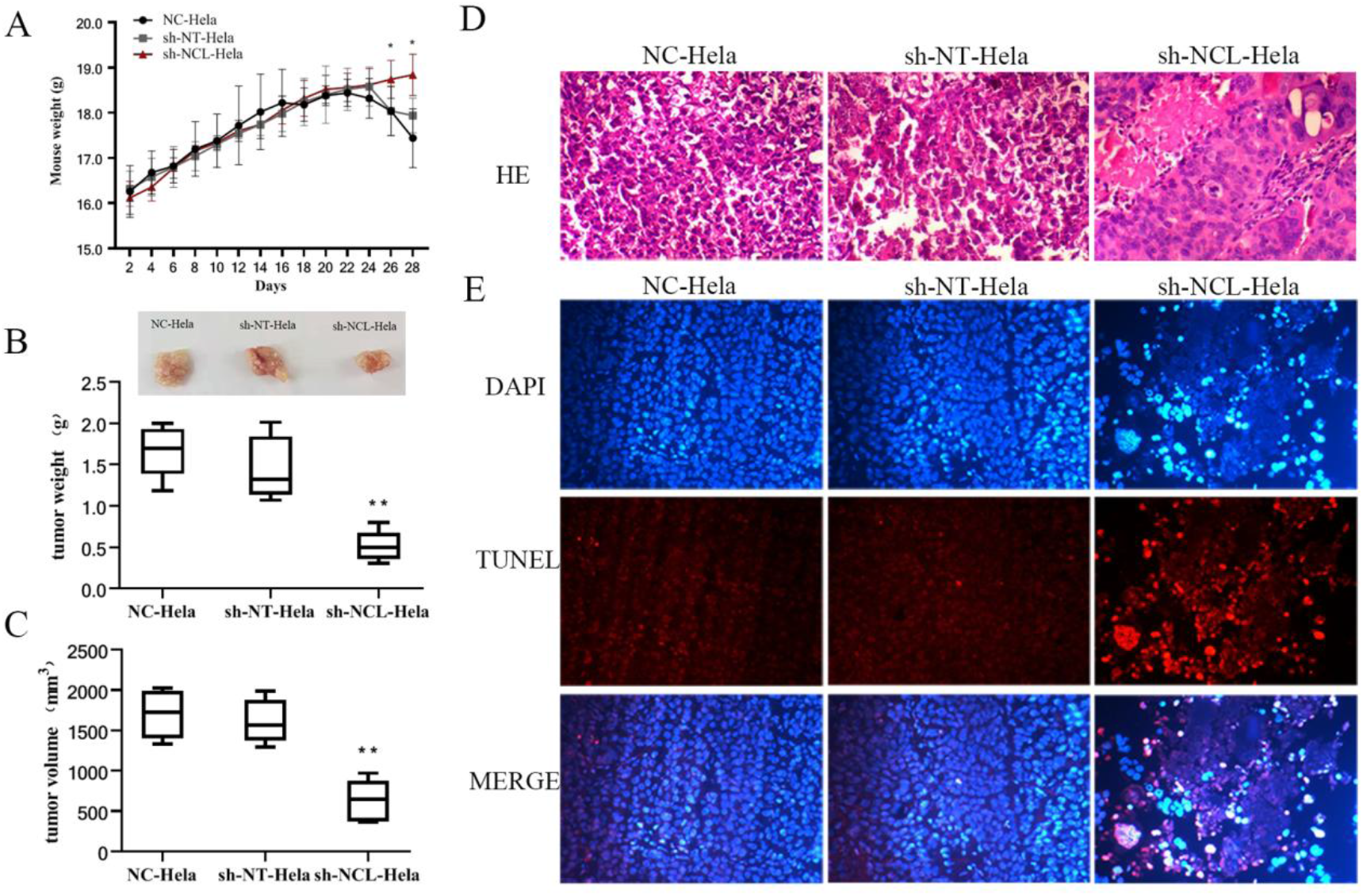
NCL knockdown regulates tumor formation in mice injected with Hela cells. **A.** The weight of each mouse was measured every two days and results were presented as a growth curve. **B.** Tumor weight was measured at the end of the day. **C.** Tumor volume was measured at the end of the day. **D.** H&E staining (×200). **E.** DAPI and TUNEL staining (×200). Results are presented as mean ± standard deviation (n = 5). ^*^*P* < 0.05, ^**^*P* < 0.01, vs. the sh-NT-Hela group, *n*=5.

### The PI3K/AKT pathway plays an important role in NCL regulated cancer cell proliferation and apoptosis

Three groups cells were collected and RNA sequencing was performed to reveal mechanisms behind induced apoptosis. Heat map analysis showed that gene expression patterns differed between the sh-NCL-Hela cells and control cell lines (sh-NT-Hela, NC-Hela, Figure 4A). A total of 2709 differentially expressed genes were identified using a fold-change cut-off of > 1.5 between sh-NCL-Hela vs sh-NT-Hela. A scatter plot of the commonly enrichment pathways was presented by KOBAS analysis (Figure 4B-C), which indicated that the number of genes in the PI3K/AKT pathway were enriched in the profiling data. In addition, the mRNA expression of several significantly altered genes was confirmed using RT-PCR (Figure 4D). Western blotting revealed that there was a significant difference in protein expression levels between sh-NT-Hela and sh-NCL-Hela cells (^*^*P*<0.01, Figure 4E), the expression levels of PI3K, P-AKT, AKT and BCL-2 in sh-NCL-Hela cells significantly decreased, while the expression levels of cleaved-Caspase 3 and Bax significantly increased.

**Figure 4:**
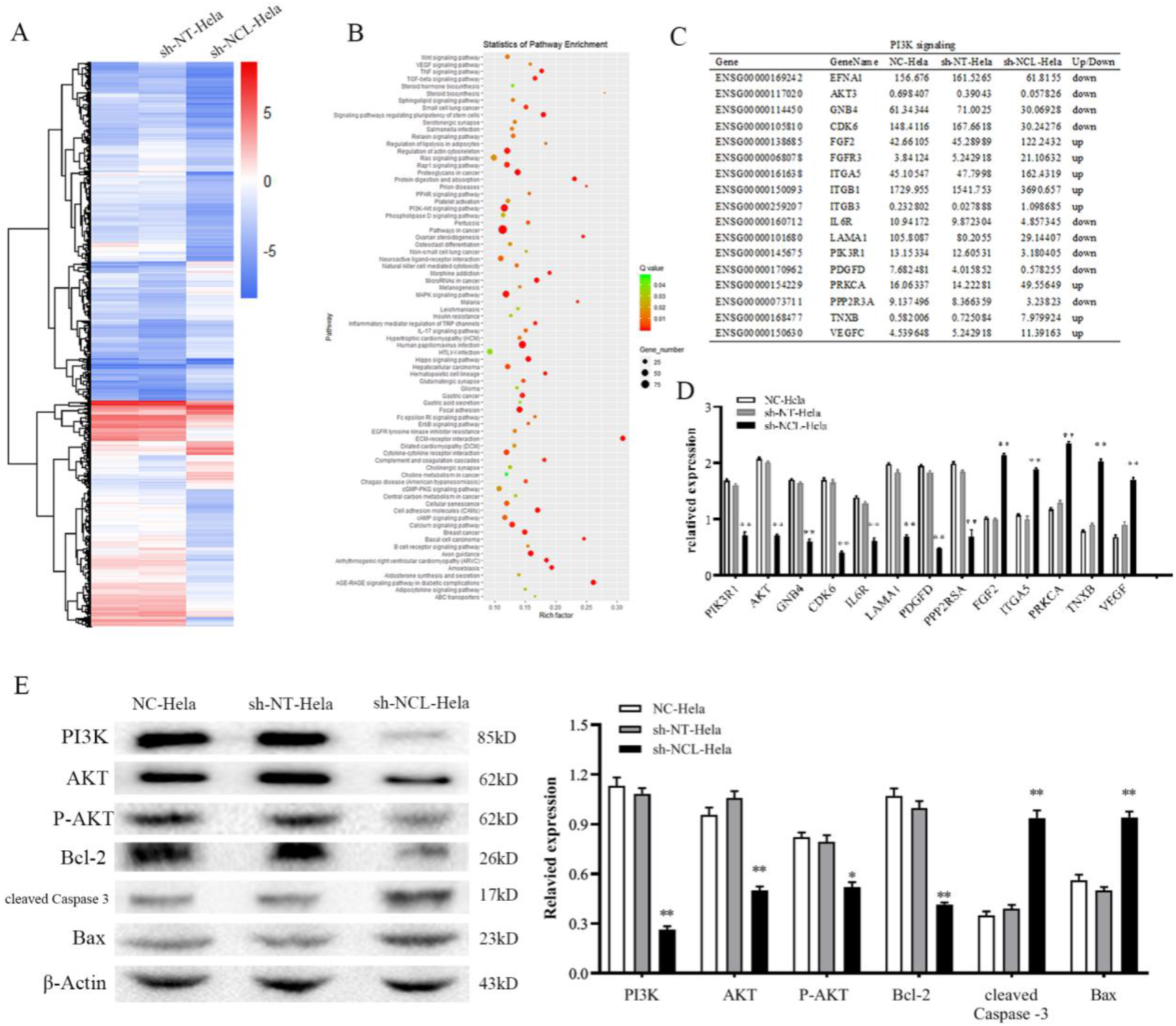
NCL-knockdown induces apoptosis by inhibiting the PI3K/AKT pathway. **A.** A heat map of differentially expressed genes in all samples. Rows represent genes whereas columns represent samples. **B.** Pathway enrichment analysis of transcriptome profiling. **C.** Enrichment of differentially expressed genes in the PI3K/AKT pathway. **D.** Diagram showing the fold-change of several typical genes using RT-PCR. **E.** Protein levels of PI3K, AKT, Bcl-2, Bax, and cleaved Caspase 3 were detected by western blotting. ^*^*P* < 0.05, ^**^*P* < 0.01, vs. the sh-NT-Hela group.

### Effects of NCL repression combined with PI3K pathway activation/ inhibition on Hela cell phenotypes

To confirm the association between NCL and the PI3K/AKT pathway, cells were treated with a PI3K inhibitor (LY294002) or a PI3K activator (740Y-P). MTT results revealed that after treatment with LY294002, the cell proliferation in each groups decreased, where the sh-NCL-Hela group showed a significantly weaker proliferation rate compared with the sh-NT-Hela group (^*^*P* < 0.05, ^#^*P*<0.05, Figure 5A). After treatment with 740Y-P, cell proliferation was enhanced, with the sh-NCL-Hela group showing significantly lower proliferation rates compared with the sh-NT-Hela group (Figure 5A). In addition, RT-PCR analyses revealed that mRNA expression levels of PI3K and AKT were lower after LY294002 treatment, with an opposite effect observed when cells were treated with 740Y-P (Figure 5B). These results were consistent with results observed for sh-NCL-Hela cells. Thus, NCL was found to play a role in inhibiting the proliferation and promoting apoptosis of Hela cells through the PI3K/AKT pathway.

**Figure 5:**
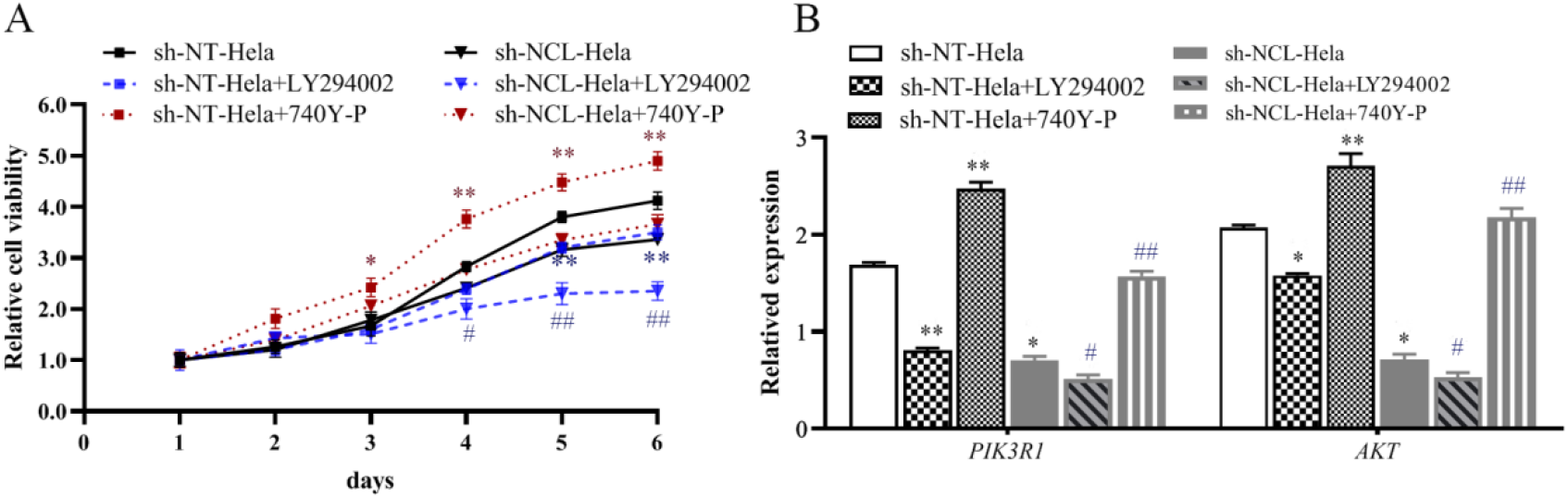
Hela cells with repressed NCL levels were treated with the PI3K inhibitor LY294002 or the PI3K activator 740Y-P. **A.** Cell proliferation was determined using a MTT assay. **B.** The expression levels of the apoptosis-related genes were measured using RT-PCR. ^*^*P* < 0.05, ^**^*P* < 0.01, vs. the sh-NT-Hela group. ^#^*P* < 0.05, ^##^*P* < 0.01, vs. the sh-NCL-Hela group, *n*=3.

## Discussion

Studying NCL in malignant tumors began in the 1980s (10) and its high expression was identified in different cancers, such as liver cancer (11), prostate cancer (12), renal cell carcinoma (13), lung cancer (14) and colon cancer (15). High expression of NCL promotes cell proliferation, angiogenesis, metastasis, and increased resistance to apoptosis in cancer cells. The distribution and structure of NCL in cells are closely related to pathological stage, tumor grade, increased disease risk and patient survival rate (16). Our results showed that knockdown of NCL inhibited the proliferation and invasion of Hela cells by promoting apoptosis. The tumor volume of nude mice inoculated with NCL knockdown cells was significantly smaller than that was observed in mice injected with normal and negative control group cell lines. In addition, tumor growth was significantly slower, indicating that NCL knockdown significantly inhibited cervical cancer growth, which was consistent with the previous findings observed in glioma (17).The results of pathological sections also revealed that downregulation of NCL expression induced apoptosis of transplanted tumor cells (Figure 3). Even though there is accumulating evidence of NCL pro-tumor activity, data are still limited and the underlying molecular mechanisms remain unclear.

Transcriptomics allow us to interpret the functional elements of the genome and reveal overall gene expression profiles associated with tumors. This field has been used in cancer research and has led to an in-depth understanding of tumorigenesis. The discovery of genes in the biological processes behind oncogenesis enhances the validation of potential diagnostic biomarkers. Ma *et al*. showed that UBC and NF-κB were TLR9 target genes associated with HPV16 infection. TLR9 may activate UBC and NF-κB through its signal transduction pathway and play a role in the occurrence and development of cervical cancer through the activation of NF-kB downstream target genes (18). Li *et al*. transcriptome analysis using Hela cells and Hela/Dox cells to find 2562 differentially expressed genes. Moreover, many key genes in these pathways were expressed differently, such as the PI3K/AKT pathway and MAPK pathway. In addition, most signaling pathways were linked to cell proliferation, signal transduction, transcription and DNA repair (19). In this study, we showed that 40 classical signaling pathways were remarkably enriched, where the PI3K/AKT pathway was the most prominent. It is also important to note that PI3K, AKT, GNB4, CDK6, IL6R, LAMA, PDGFD, PPP2RSA, FGF, ITGA, PRKCA, TNXB, VEGF and other genes in the pathway were all significantly altered.

Previous studies demonstrated that signal transmission in the PI3K/AKT pathway mainly relies on phosphorylation level changes by upstream key molecules to activate downstream protein factors. This pathway is then involved in transmitting signals from different cytokines to the nucleus and has a close and complex relationship with biological processes, such as cell survival, proliferation, apoptosis, invasion and metastasis (20). Serine/threonine kinase AKT, the most important mediator, plays a major role in signal transduction throughout the PI3K pathway. Consequently, AKT was selected and tested for changes in transcription and expression levels regulated by NCL. This study revealed that the downregulation of NCL reduced AKT expression and inhibited PI3K/AKT pathway. Further data showed that GβY, Cytokine R, LAMA, and PDGFD, all of which are downstream genes of PI3K/AKT, were downregulated, whereas FGF, ITGA, TNXB and VEGF genes were up-regulated in the sh-NCL-Hela cells, which exerted their effects on CDK6, PRKCA and PKCs. Cyclin-dependent kinase 6 (CDK6) plays a vital role in the regulation of cell cycle progression. It is the catalytic subunit of the CDK6-cyclin D complex, which participates in progression from G1 to S phases of the cell cycle and negatively regulates cell differentiation. Its activity first appears in the middle of the G1 phase to regulate the activity of tumor suppressor protein retinoblastoma (Rb). Furthermore, CDK6 can promote the proliferation of tumor cells and is dysregulated in breast cancer, lymphoma, glioma and melanoma (21). Overexpression of CDK6 reduces skin tumorigenesis and the growth of fibroblasts. MicroRNAs (miRNAs) blocking CDK6 inhibit the proliferation of glioma and lung cancer (22). These observations, combined with the fact that mice lacking D-type cyclins and/or CDK4/6 are viable, make the cyclin D-INK4-pRb-E2F pathway of cyclins an attractive anticancer target (23). Protein kinase Cα (PRKCA) is a serine/ threonine protein kinase and a member of the PKC family. The PKC family regulates the ERK-MAPK, PI3K/AKT and NF-κB pathways and is involved in a variety of cellular functions, such as cell proliferation and survival (24). PRKCA is upregulated in several human cancer types, including breast cancer (25), non-small cell lung cancer (26) and hematological malignancies (27). Protein phosphatase 2 regulatory subunit B alpha (PPP2R3A), is regulates several important signal transduction pathways related to cancer, including the Wnt signaling cascade, adenosine monophosphate activated protein kinase (AMPK) activity, and epidermal growth factor (EGF)/EGF receptor (EGFR) signaling pathway. PP2A is known to be involved in tumorigenesis and its function balances kinase-mediated phosphorylation throughout cell signaling networks. Previous studies have shown that PP2A phosphorylates Akt and ERK. One report showed that treatment with a okadaic acid (OA), a PP2A inhibitor, not only inhibited PP2A activation but also triggered Akt and ERK signal transduction and increased cell growth, migration, and angiogenic ability. Thus, this supported the anti-proliferation effects of PP2A in endothelial cells. Zhou *et al*. found that overexpression of bcl-2 inhibited caspase-3 activity, while caspase-3 inactivation alleviated PP2A degradation, leading to a dissociation between PP2A and Akt (28). Caspase-3 is the main executor caspase that can be cleaved and activated bycaspase-8, caspase-9 and caspase-10. Lysis of p12 and p17 subunits at aspartate residues form active caspase-3 enzyme that degrades a variety of cellular proteins and leads to morphological changes and DNA fragmentation during apoptosis (29).Mechanistically, our data from knockout studies indicated that the expression levels of active caspase-3 was higher in sh-NCL-Hela cells than in sh-NT-Hela cells and was associated with apoptosis. At the same time, Akt phosphorylation of Bax changes it from being pro- to anti-apoptotic, resulting in reduced mitochondrial sensitivity to apoptotic signal transduction. These reports both confirm and extend our results that the effects of NCL were related to the activation of PI3K/AKT pathways in Hela cells. To assess the relationship between NCL and the PI3K/AKT pathway in cervical cancer, activators and inhibitors of the PI3K/AKT pathway were used to observe changes in the two groups. The expression levels of apoptosis-related genes in Hela cells after using PI3K inhibitor LY294002 were consistent with sh-NCL-Hela cells, while the effects of the agonist 740Y-P were opposite. Altogether, these data indicate that decreased NCL expression may promote cell apoptosis by an independent mechanism in part by the PI3K/AKT pathway.

In conclusion, NCL plays an important role in the occurrence and development of cervical cancer and may be used as a biological marker for cervical cancer prevention and treatment. Further research is needed to elucidate the potential mechanism of other signaling pathways involved in the antiapoptotic effects of NCL and the changes of specific sites and domains in the PI3K/AKT signaling pathway.

## Experimental Procedures

### Cell lines

Human Hela cervical cancer cells were obtained from the Cell Bank of Chinese Academy of Sciences and cultured in DMEM (Thermo Fisher Scientific, Waltham, USA) supplemented with 10% fetal bovine serum (FBS; Sijiqing, Hangzhou, China), 2mM L-glutamine and antibiotics (100 U/mL streptomycin and 100 U/mL penicillin). Cells were maintained in a 37°C humidified 5% CO_2_ incubator.

### Lentivirus constructs and transfections

Cells were designated into a blank control group (NC-Hela, uninfected Hela cells), negative control group (sh-NT-Hela, infected with negative control vector, target sequence: 5’-TTCTCCGAACGTGTCACGT-3’) or a NCL experimental group (sh-NCL-Hela, infected with NCL knockdown vector, target sequence: 5’-GGAAATGTCAGAAGATGAAGA-3’). The RNAi lentiviral vector packaging system was constructed by Genechem (Shanghai, China). Hela cells were transduced using a lentivirus expressing shRNA when they reached 70% confluency in serum-free growth medium under a multiple infection (MOI) of 20 particles/cell. Three days after transduction, GFP expression was observed using fluorescence microscopy and puromycin was selected to establish a stable cell line. Infection efficiency was determined using RT-PCR and western blotting assays (30).

### RNA isolation and quantification

Total RNA in different treatment groups was obtained using TRIzol^®^ reagent (Invitrogen™, Carlsbad, CA, USA) according to instructions provided by the manufacturer and 2μg aliquots were reversed transcribed cDNAs using PrimeScript™ RT-PCR Kit (Takara, Dalian, China). After, cDNAs were subjected to PCR analyses using specific primers (The primer sequences are listed in Table 1) on an amplification detection system (Bio-Rad, Hercules, USA). β-actin served as a housekeeping gene. Messenger RNA (mRNA) expression levels were expressed as the ratio of the gray level of each sample to its internal reference β-actin control and analyzed by QuantityOne program (Bio-Rad, Hercules, USA) (31).

**TABLE 1.**
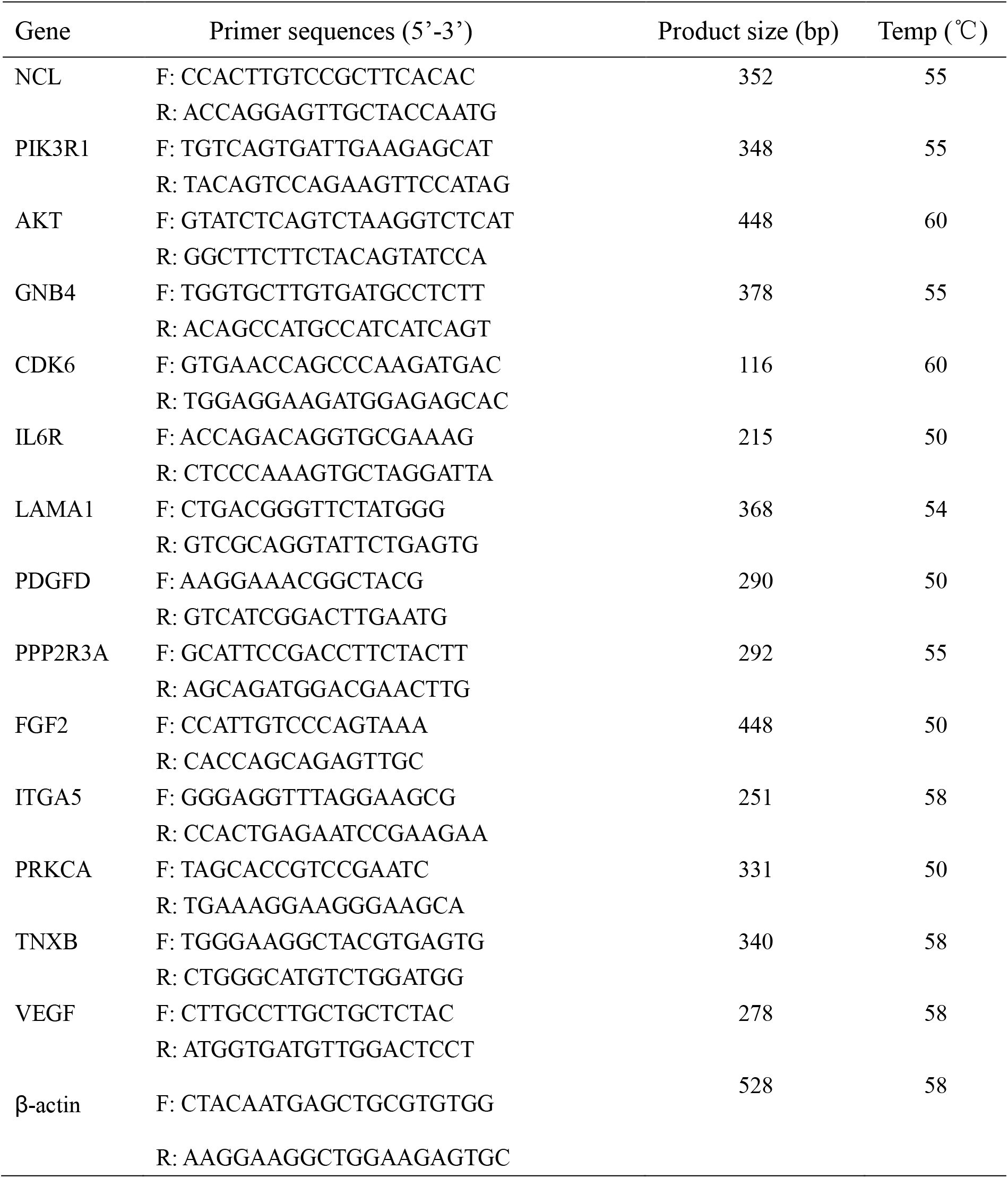
Primer sequences and product sizes

### Western blotting analysis

Protein was extracted from cells using RIPA lysis buffer and protein levels were measured using a BCA kit (Beyotime, Shanghai, China). Then, 30 μg protein samples were separated using a 10% SDS-PAGE and transferred to a PVDF membrane. Next, the membrane was washed using 5% skimmed milk diluted in TBST for 2 h, followed by an incubation with primary antibodies at 4 °C overnight. After being washed with TBST (containing 0.1% Tween-20), the membranes were incubated with corresponding secondary antibodies for 2 h at room temperature. Immune complexes were examined using ECL detection (Beyo ECL Plus; Beyotime, Shanghai, China). Protein bands were quantified using ImageJ Plus software. The primary antibodies used in western blotting included: anti-NCL (No.10556), anti-PI3K (No. 60225), anti-AKT (No. 60203), anti-Phospho-AKT (No. 66444), anti-Bax (No. 50599), anti-Caspase-3 (No.19677), anti-BCL-2 (No. 12789) and anti-β-actin (No. 20536). These antibodies were purchased from Proteintech Group, Inc. (Chicago, USA) (32).

### Cell proliferation assays

Cells (1000 per well) were seeded into 96-well plates and incubated at 37 °C for 1 to 6 days. Next, 3-(4, 5-Dimethylthiazol-2-yl)-2, 5-diphenyltetrazolium bromide (MTT) was added to each well at a final concentration of 0.5 mg/mL for 4 h. The supernatants were transferred, dimethyl sulfoxide (DMSO) was added and the absorbance of each well was measured at 490 nm using an ELISA microplate reader (Bio-Rad, Hercules, CA, USA) (33).

### Colony formation assays

Cells (density of 5 × 10^2^ per well) were re-seeded into 6-well plates and incubated for 7 to 14 days at 37 °C, with medium changed every 3 days. Cells were fixed in 100% methanol for 15 min at 4 °C and then stained with crystal violet (1%, wt/vol; Beyotime, Shanghai, China) for 15 min. An inverted microscope was used to count the number of newly formed colonies > 200 mm in diameter (34).

### HOECHST 33342 staining

Cells in each group that were cultured for 48 h after subculture were washed twice with PBS. Next, 1 mL of 5 μg/mL Hoechst 33342 (Beyotime, Shanghai, China) working solution was added and cells were incubated for 30 min at 37 °C, observed and photographed using an inverted fluorescence microscope (35).

### Detection of apoptosis by FCM

Annexin V-APC/7AAD Apoptosis Detection kit (BD Biosciences, San Jose, CA, USA) was used to measure apoptosis levels, following manufacturer instructions. Cells were collected, washed twice with cold PBS and gently rocked on a shaker in 100 μL 1 × binding buffer. The binding buffer contained 2.5 μL of APC coupled annexin-V and 5.0 μL of 7-AAD. The buffer was then incubated for 15 min in the dark at room temperature. Stained cells were analyzed using flow cytometry (36).

### Caspase activity assay

Caspase −3 Activity Assay Kit (Beyotime, Shanghai, China) was used to detect Caspase −3 enzyme activity in cells by a colormetric assay. Cells were washed twice with PBS, and then lysed by 150 μL lysis buffer on ice for 20 minutes. After being centrifuged at 12000 rpm at 4 °C for 15 min, the supernatants were collected and quantified. 50 μL of supernatants (protein concentration reaches 1-3mg/mL) was mixed with 40 μL buffer and 10 μL of Caspase reaction substrate (2mM Ac-DEVD-pNA), incubated at 37 °C for 1.5 h. The relative activity of Caspase was determined with a spectrophotometer (DU-800, BECKMAN, USA) at 405 nm (31).

### In vivo tumorigenic assays

BALB/C-nu mice aged 4-6 weeks were purchased from the Shanghai SLAC Laboratory Animal Center of the Chinese Academy of Sciences (Shanghai, China). All animals were raised and maintained under pathogen free conditions. All experiments were approved by the Animal Conservation and Use Committee and were performed following institutional guidelines. Each group of cells (1 × 10^7^ cells in 200 μL PBS) was subcutaneously injected into the groin area of anaesthetized nude mice. Tumor formation was recorded by measuring the maximum diameter of the length (a) and width (b) every two days. The following formula was used to calculate the tumor volume: V (mm^3^) = 0.5 × a × b^2^. Four weeks later, nude mice were euthanized using carbon dioxide and the final volume and weight of the tumor were recorded. Each tumor was fixed in 4% paraformaldehyde for 48 h, embedded in paraffin and sliced into 4 μm tissue sections for hematoxylin and eosin (H&E) staining (37) and TUNEL staining (38). A one-step TUNEL apoptosis detection kit (KeyGen Biotech, Jiangsu, China) was used to detect apoptotic cells in tumor tissues. In addition, 4’,6-diamidino-2-phenylindole (DAPI) was used to stain nuclei. Digital images were captured using a fluorescence microscope (Nikon, Tokyo, Japan).

### Gene expression profiling

A total of 2 × 10^5^ Hela cells were inoculated into 6-well plates for 48 h. Each sample was repeated three times to extract RNA. PureLink™ RNA Mini Kit (ThermoFisher Scientific) was used to separated total RNA based on instructions provided by the manufacturer. A Qubit ^®^ 3.0 Flurometer (Life Technologics, CA, USA) was used to identify the quantity and quality of RNA samples to ensure that all samples RIN > 7.0. RNA samples were sent to Anoroad (Beijing, China) for library construction and sequencing. An Agilent 2100 RNA Library Prep Kit for Illumina (Agilent Technologies, CA, USA) was used to prepare the RNA library. Trim_galore (v0.4.2, Babraham Bioinformatics) was used to trim the low-quality and cohesive sequences in the original reads and RNA-STAR (v2.4.0j) was used to map the clean reads to the reference genome (human genome 19). DEGSeq was used for genetic difference analysis, compared with the treatment group and the reference group and differential genes were selected with |log_2_Ratio|≥ 1.5 and q < 0.05. KOBAS analysis of Gene Ontology (GO) and Kyoto Encyclopedia of Genes and Genomes (KEGG) for DE genes were used. The R Pathview software package was used to visualize gene expression status in KEGG pathways (39).

### Statistical Analysis

All statistics were analyzed using SPSS 19.0 (SPSS Standard version 19.0, SPSS Inc., Chicago, IL). A student’s *t* test and one-way analysis of variance (ANOVA) were performed to compare different groups. Data were presented as the mean ± standard deviation (SD). All data were from at least three independent experiments. A *P* value < 0.05 was considered statistically significant, with the following notations: ^*^*P* < 0.05; ^**^*P* < 0.01.

## Data Availability Statement

The data that support the findings of this study are available from the corresponding author upon reasonable request.

## Acknowledgements

This research was funded by Zhejiang Provincial Natural Science Foundation of China under Grant No. LQ17H160015; Zhejiang Province welfare technology applied research project under Grant No. LGC20H260001; National Natural Science Foundation of China under Grant No.81501808; the Wenzhou City Science and Technology Foundation of China under Grant No. Y20190052 and Y20170032; the National Undergraduate Training Program for Innovation and Entrepreneurship of China under Grant No. 202010343022.

## Author Contributions

Conceptualization, J.Y., Q.B. and P.L.; methodology and investigation, R.P., Z.T., J.Z., P.R., Y.L., E.Z., D.H., and P.H.; funding acquisition and review, D.L., Q.B. and P.L.; writing-review and editing, P.L. and J.Y.; project administration, P.L.. All authors have read and agreed to the pulished version for the manuscript.

## Institutional Review Board Statement

All the experiments involving in animals were reviewed and approved by The Ethics Committee of Wenzhou Medical University.

## Informed Consent Statement

Not applicable for studies not involving humans.

## References

1. Zhou, C., Li, C., Peng, S., Zhou, L., and Li, H. (2020) Comprehensive Analysis of the Relationships Between Tumor Mutation Burden With Immune Infiltrates in Cervical Cell Carcinoma. Front.Mol. Biosci. 7, 582911.

2. Cheng, HR., and Chen, J. (2016) Advances in diagnosis and treatment of early stage cervical cancer in young patients. Journal of Modern Oncology. 24, 24678–680.

3. Feng, R.M., Zong, Y.N., Cao, S.M., and Xu, R.H. (2019) Current cancer situation in China: good or bad news from the 2018 Global Cancer Statistics? Cancer. Commun (Lond). 39, 22.

4. Ugrinova, I., Monier, K., Ivaldi, C., Thiry, M., Storck, S., Mongelard, F., and Bouvet, P. (2007) Inactivation of nucleolin leads to nucleolar disruption, cell cycle arrest and defects in centrosome duplication. BMC. Mol. Biol. 8, 66.

5. Reyes-Reyes, E. M., and Akiyama, S. K. (2008) Cell-surface nucleolin is a signal transducing P-selectin binding protein for human colon carcinoma cells. Exp. Cell. Res. 314, 2212–2223.

6. Otake, Y., Soundararajan, S., Sengupta, T. K., Kio, E. A., Smith, J. C., Pineda-Roman, M., Stuart, R. K., Spicer, E. K., and Fernandes, D. J. (2007) Overexpression of nucleolin in chronic lymphocytic leukemia cells induces stabilization of bcl2 mRNA. Blood. 109, 3069–3075.

7. Morimoto, Y., Kito, S., Ohba, T., Morimoto, H., Okamura, H., and Haneji, T. (2001) Alteration of argyrophilic nucleolar organizer region associated (Ag-NOR) proteins in apoptosis-induced human salivary gland cells and human oral squamous carcinoma cells. J. Oral. Pathol. Med. 30, 193–199.

8. Kito, S., Shimizu, K., Okamura, H., Yoshida, K., Morimoto, H., Fujita, M., Morimoto, Y., Ohba, T., and Haneji, T. (2003) Cleavage of nucleolin and argyrophilic nucleolar organizer region associated proteins in apoptosis-induced cells. Biochem. Biophys. Res. Commun. 300, 950–956.

9. Sanjeev, K., Elizabhet, C. G., Mounira, C.D., Cong, R., Sadhan, D., Iva, U., Xavier, G., Karine, M., Fabien, M., and Philippe, B. (2017) Integrated analysis of mRNA and miRNA expression in HeLa cells expressing low levels of Nucleolin. Sci. Rep. 7, 9017.

10. Orrick, L. R., Olson, M. O., and Busch, H. (1973) Comparison of nucleolar proteins of normal rat liver and Novikoff hepatoma ascites cells by two-dimensional polyacrylamide gel electrophoresis. Proc. Natl. Acad. Sci. USA. 70, 1316–1320.

11. Chen, S.C., Hu, T.H., Huang, C.C., Kung, M.L., Chu, T.H., Yi, L.N., Huang, S.T., Chan, H.H., Chuang, J.H., Liu, L.F., Wu, H.C., Wu, D.C., Chang, M.C., and Tai, M.H. (2015) Hepatoma-derived growth factor/ nucleolin axis as a novel oncogenic pathway in liver carcinogenesis. Oncotarget. 6, 16253–16270.

12. Masiuk, M., Lewandowska, M., Dobak, E., and Urasinska, E. (2020) Nucleolin and Nucleophosmin Expression in Gleason 3 and Gleason 4 Prostate Cancer With Seminal Vesicles Invasion (pT3b). Anticancer. Res. 40, 1973–1979.

13. Rosenberg, J. E., Bambury, R. M., Van Allen, E. M., Drabkin, H. A., Lara, P. N., Harzstark, A. L., Wagle, N., Figlin, R. A., Smith, G. W., Garraway, L. A., Choueiri, T., Erlandsson, F., and Laber, D. A. (2014) A phase II trial of AS1411 (a novel nucleolin-targeted DNA aptamer) in metastatic renal cell carcinoma. Invest. New. Drugs. 32, 178–187.

14. Huang, F., Wu, Y., Tan, H., Guo, T., Zhang, K., Li, D., and Tong, Z. (2019) Phosphorylation of nucleolin is indispensable to its involvement in the proliferation and migration of non-small cell lung cancer cells. Oncol. Rep. 41, 590–598.

15. Satake, Y., Kuwano, Y., Nishikawa, T., Fujita, K., Saijo, S., Itai, M., Tanaka, H., Nishida, K., and Rokutan, K. (2018) Nucleolin facilitates nuclear retention of an ultraconserved region containing and accelerates colon cancer cell growth. Oncotarget. 9, 26817–26833.

16. Berger, C. M., Gaume, X., and Bouvet, P. (2015) The roles of nucleolin subcellular localization in cancer. Biochimie. 113, 78–85.

17. Xu, Z., Joshi, N., Agarwal, A., Dahiya, S., Bittner, P., Smith, E., Taylor, S., Piwnica-Worms, D., Weber, J. and Leonard, J. R. (2012) Knocking down nucleolin expression in gliomas inhibits tumor growth and induces cell cycle arrest. J. Neurooncol. 108, 59–67.

18. Ma, ML., Tuerxun, MLB., Wang, ZF., and Axianggu, H. (2018) Effect of HPV16 infection on the expression of UBC AND NF-κB in cervical lesions based on transcriptome sequencing. Journal of Xinjiang Medical University. 41, 534–538.

19. Li, PK. (2018) Study on acquired drug resistance of human cervical cancer Hela cells doxorubicin based on transcriptome sequencing. Xian: Northwest University.

20. Fresno Vara, J. A., Casado, E., de Castro, J., Cejas, P., Belda-Iniesta, C., and González-Barón, M. (2004) PI3K/Akt signalling pathway and cancer. Cancer. Treat. Rev. 30, 193–204.

21. Zhu, K., Liu, L., Zhang, J., Wang, Y., Liang, H., Fan, G., Jiang, Z., Zhang, C.Y., Chen, X., and Zhou, G. (2016) MiR-29b suppresses the proliferation and migration of osteosarcoma cells by targeting CDK6. Protein. Cell. 7, 434–444.

22. Li, B., He, H., Tao, B.B., Zhao, Z.Y., Hu, G.H., Luo, C., Chen, J.X., Ding, X.H., Sheng, P., Dong, Y., Zhang, L., and Lu, Y.C. (2012) Knockdown of CDK6 enhances glioma sensitivity to chemotherapy. Oncol. Rep. 28, 909–914.

23. Hamilton, E., and Infante, J. R. (2016) Targeting CDK4/6 in patients with cancer. Cancer Treat Rev. 45, 129–138.

24. Fu, Q., Song, X., Liu, Z., Deng, X., Luo, R., Ge, C., Li, R., Li, Z., Zhao, M., Chen, Y., Lin, X., Zhang, Q., and Fang, W. (2017) miRomics and Proteomics Reveal a miR-296-3p/PRKCA/FAK/Ras/c-Myc Feedback Loop Modulated by HDGF/DDX5/β-catenin Complex in Lung Adenocarcinoma. Clin. Cancer. Res. 23, 6336–6350.

25. Lahn, M., Köhler, G., Sundell, K., Su, C., Li, S., Paterson, B. M., and Bumol, T. F. (2004) Protein kinase C alpha expression in breast and ovarian cancer. Oncology. 67, 1–10.

26. Arora, S., Ranade, A. R., Tran, N. L., Nasser, S., Sridhar, S., Korn, R. L., Ross, J. T. D., Dhruv, H., Foss, K. M., Sibenaller, Z., Ryken, T., Gotway, M. B., Kim, S., and Weiss, G. J. (2011) MicroRNA-328 is associated with (non-small) cell lung cancer (NSCLC) brain metastasis and mediates NSCLC migration. Int. J. Cancer. 129, 2621–2631.

27. Saloustros, E., Salpea, P., Qi, C.-F., Gugliotti, L. A., Tsang, K., Liu, S., Starost, M. F., Morse, H. C., and Stratakis, C. A. (2015) Hematopoietic neoplasms in Prkar2a-deficient mice. J. Exp. Clin. Cancer. Res. 34, 143.

28. Zhou, H., Luo, W., Zeng, C., Zhang, Y., Wang, L., Yao, W., and Nie, C. (2017) PP2A mediates apoptosis or autophagic cell death in multiple myeloma cell lines. Oncotarget. 8, 80770–80789.

29. Enari, M., Sakahira, H., Yokoyama, H., Okawa, K., Iwamatsu, A., and Nagata, S. (1998) A caspase-activated DNase that degrades DNA during apoptosis, and its inhibitor ICAD. Nature. 391, 43–50.

30. Xu, Z., Zhu, C., Chen, C., Zong, Y., Feng, H., Liu, D., Feng, W., Zhao, J., and Lu, A. (2018) CCL19 suppresses angiogenesis through promoting miR-206 and inhibiting Met/ERK/Elk-1/HIF-1α/VEGF-A pathway in colorectal cancer. Cell. Death. Dis. 9, 974.

31. Pan, R., Lu, R., Zhang, Y., Zhu, M., Zhu, W., Yang, R., Zhang, E., Ying, J., Xu, T., Yi, H., Li, J., Shi, M., Zhou, L., Xu, Z., Li, P., and Bao, Q. (2015) Spirulina phycocyanin induces differential protein expression and apoptosis in SKOV-3 cells. Int. J. Biol. Macromol. 81, 951–959.

32. Wu, L.C., Lin, Y.Y., Yang, S.Y., Weng, Y.T., and Tsai, Y.T. (2011) Antimelanogenic effect of c-phycocyanin through modulation of tyrosinase expression by upregulation of ERK and downregulation of p38 MAPK signaling pathways. J. Biomed. Sci. 18, 74.

33. Kim, J.Y., Jung, E. J., Park, H. J., Lee, J.H., Song, E. J., Kwag, S.J., Park, J.H., Park, T., Jeong, S.H., Jeong, C.Y., Ju, Y.T., Lee, Y.J., and Hong, S.C. (2018) Tumor-Suppressing Effect of Silencing of Annexin A3 Expression in Breast Cancer. Clin. Breast. Cancer. 18, e713–e719.

34. Xu, Q., Wu, N., Li, X., Guo, C., Li, C., Jiang, B., Wang, H., and Shi, D. (2019) Inhibition of PTP1B blocks pancreatic cancer progression by targeting the PKM2/AMPK/mTOC1 pathway. Cell. Death. Dis. 10, 874.

35. Crowley, L. C., Marfell, B. J., and Waterhouse, N. J. (2016) Analyzing Cell Death by Nuclear Staining with Hoechst 33342. Cold. Spring. Harb. Protoc. 2016, 9.

36. Pan, H., Pan, J., Ji, L., Song, S., Lv, H., Yang, Z., and Guo, Y. (2019) Carboxypeptidase A4 promotes cell growth via activating STAT3 and ERK signaling pathways and predicts a poor prognosis in colorectal cancer. Int. J. Biol. Macromol. 138, 125–134.

37. Heidari, M., Heidari-Vala, H., Sadeghi, M. R., and Akhondi, M. M. (2012) The inductive effects of Centella asiatica on rat spermatogenic cell apoptosis in vivo. J. Nat. Med. 66, 271–278.

38. Lin, P., Cheng, Y., Song, S., Qiu, J., Yi, L., Cao, Z., Li, J., Cheng, S., and Wang, J. (2019) Viral Nonstructural Protein 1 Induces Mitochondrion-Mediated Apoptosis in Mink Enteritis Virus Infection. J. Virol. 93, e01249–19.

39. Zhang, J., Qu, Z., Yao, H., Sun, L., Harata-Lee, Y., Cui, J., Aung, T. N., Liu, X., You, R., Wang, W., Hai, L., Adelson, D. L., and Lin, L. (2019) An effective drug sensitizing agent increases gefitinib treatment by down regulating PI3K/Akt/mTOR pathway and up regulating autophagy in non-small cell lung cancer. Biomed. Pharmacother. 118, 109169.

